# Optimum time for hand pollination in yam (*Dioscorea* spp.)

**DOI:** 10.1101/2022.05.26.493634

**Authors:** Jean M. Mondo, Paterne A. Agre, Robert Asiedu, Malachy O. Akoroda, Asrat Asfaw

## Abstract

Hand pollination success rate is low in yam *(Dioscorea* spp.), due partly to suboptimal weather conditions. Thus, determining the most suitable time for pollination could improve the pollination success in yam breeding programs. We performed continuous hand pollination within flowering windows of *D. rotundata* and *D. alata* for two consecutive years to determine the most appropriate month, week, and hours of the day allowing maximum pollination success. In *D. alata* crossing block, we observed significant differences among crossing hours for pollination success (p = 0.003); morning hours (8–12 a.m.) being more conducive than afternoons (12–5 p.m.). No significant differences existed between crossing hours in *D. rotundata,* though the mid-day seemed optimal. For both species, the time interval 11–12 a.m. was more appropriate for crossing while 4–5 p.m. was the poorest. However, *in vitro* pollen germination tests showed that mid-day pollen collection (12 noon – 2 p.m.) had better results than both extremes, though there were strong genotypic effects on outcomes. Pollination success rates differed significantly among months for *D. alata* (p < 0.001) but not for *D. rotundata* (p > 0.05). Differences in pollination success existed across weeks within flowering windows of both *D. alata* (p < 0.001) and *D. rotundata* (p = 0.004). The seed production efficiency (SPE) had a similar trend as the pollination success rate. No clear pattern existed between the pollination time and the seed setting rate (SSR) or seed viability (SV), though their dynamics varied with weeks and months. This study provided an insight on the dynamics of pollination outcomes under the influence of pollination times and allows detecting months, weeks, and hours of the day when hybridization activities should be focused for better results.

## Introduction

Yam (*Dioscorea* spp.) is a multispecies staple crop with significant contributions to food security and poverty alleviation in tropics and subtropics, especially in West Africa where it is extensively produced (Asiedu and Sartie, 2010). Its cultivation faces several yield restricting and quality reducing factors related to poor crop husbandry, biotic and abiotic stresses, and postharvest losses that widen the gap between the farmer yields (~10 t ha^−1^) and the crop potential (40–50 t ha^−1^) and reduce the market penetration (Frossard et al., 2017). Plant breeding research is an integral component of addressing these challenges by development and delivery of resilient, productive, and high-quality varieties. However, improved cultivar development through breeding in yam is challenged by sexual reproduction abnormalities resulting from sparse, irregular, and asynchronous flowering, cross compatibility barriers as the vegetative propagation is favored at the expense of botanical seeds during the domestication and subsequent cultivation process (Mondo et al., 2020). Yam plant exhibits extremely low levels of fruit-to-flower and seed-to-ovule ratios, partly because of the sensitivity of its reproductive phases to suboptimal weather conditions (Mondo et al., 2022). Important climatic factors such as temperature, rainfall, relative humidity, and light intensity fluctuate from one location to the other and even from time to time in the same location. These fluctuations dictate the limit within which controlled pollination can be successfully conducted in any given location for a given species (Salami, 2016; Maity et al., 2019; Raina and Kaul, 2019; Katano et al., 2020; Mondo et al., 2022).

Several attempts were undertaken to improve hand pollination success rate in yam breeding programs by determining the appropriate time for crossing (Akoroda et al., 1981; Akoroda, 1983; Abraham and Nair, 1990; Bakayoko et al., 2019; Mondo et al., 2020). However, the optimum time for yam pollination is location-specific and depends on local environmental conditions (Mondo et al., 2020). High relative humidity, well-distributed rainfall, sunshine, and moderate atmospheric temperatures are the leading climatic factors for successful pollination in *D. alata* and *D. rotundata* yams (Abraham and Nair, 1990; Mondo et al., 2022). Recommended time for hand pollination in Nigeria (12 noon–3 p.m.) was set ~40 years ago (Akoroda, 1983), thus, there is a chance that trends recorded four decades ago may have changed. Besides, due to predominant sunny conditions at the previously recommended crossing hours, crossing activities are seldom undertaken at mid-day (Mondo et al., 2020). Pollinators most conveniently operate in morning hours (8 a.m.–12 noon). Yet, no study assessed the pollination success rates at those hours compared to the mid-day hours recommended by the literature.

Most yam species, including *D. alata* and *D. rotundata*, are dioecious with male and female flowers on separate individuals (Agre et al., 2020; Mondo et al., 2020; 2021a; 2022). The gene flow between and among these species to meet breeding objectives depends, therefore, on crosspollination success. The cross-pollination involves three phases: the release of pollen from the anther, transfer of pollen from the anther to the stigma, and successful placement of pollen on receptive stigma surface, followed by germination (Di-Giovanni and Kevan, 1991; Bhattacharya and Mandal, 2004; He et al., 2017). The transfer of yam pollen from the anther to the stigma is either by the assistance of local insects (natural) or human hand (artificial) since the sticky nature of yam pollen renders the wind pollination impossible (Mondo et al., 2020). However, the insects’ inefficiency is a major factor of low natural pollination success in yam (Akoroda, 1985; Segnou et al., 1992). This insects’ inefficiency is associated with low visitation rate, limited movements, and selectivity (Martin et al., 1963; Mondo et al., 2020). Hand pollination is used as an alternative solution; it is 2–3 times more efficient than natural pollination by insects (Akoroda, 1983; Segnou et al., 1992). Whether natural or artificial, the pollination success is associated with other factors such as pollen viability, stigma receptivity, cross compatibility, and the prevailing weather conditions (Lebot et al., 2019; Mondo et al., 2020; 2022).

This study aims at improving pollination success in yam breeding programs by assessing the optimum time of pollination, when the pollen is fully viable, the stigma receptive, and the weather is conducive in *D. alata* and *D. rotundata* crossing blocks. It uses crossing block, *in vitro* pollen germination, and weather data for assessment.

## Materials and methods

### Study site, plant material, and field establishment

Two-year experiment was conducted at the International Institute of Tropical Agriculture (IITA) Ibadan (7°29’ N and 3°54’ E), Nigeria, from April 2020 to February 2022. Six female parents (three *D. rotundata* and three *D. alata*) were selected based on the length of their flowering window and flowering intensity. On the other hand, six male parents (three per species) were used as pollen sources. All these materials were breeding lines maintained by the IITA Yam Breeding Unit (S1 Table). The cross-compatibility among selected genotypes and their ploidy statuses were based on historical data information (Mondo et al., 2022).

The planting was done in April for both species and seasons. Male and female crossing blocks were grown at appropriate spacing (1 m × 1 m). Recommended field management was conducted, including individual plant staking, fertilizer application, supplemental irrigation, regular weeding, etc. Pollination on *D. rotundata* crossing blocks were carried out from August to mid-October while it started in late September and ended in early December for *D. alata.* Weather data in the field was recorded using a data logger for the entire research period (Fig 1).

**Figure 1.**
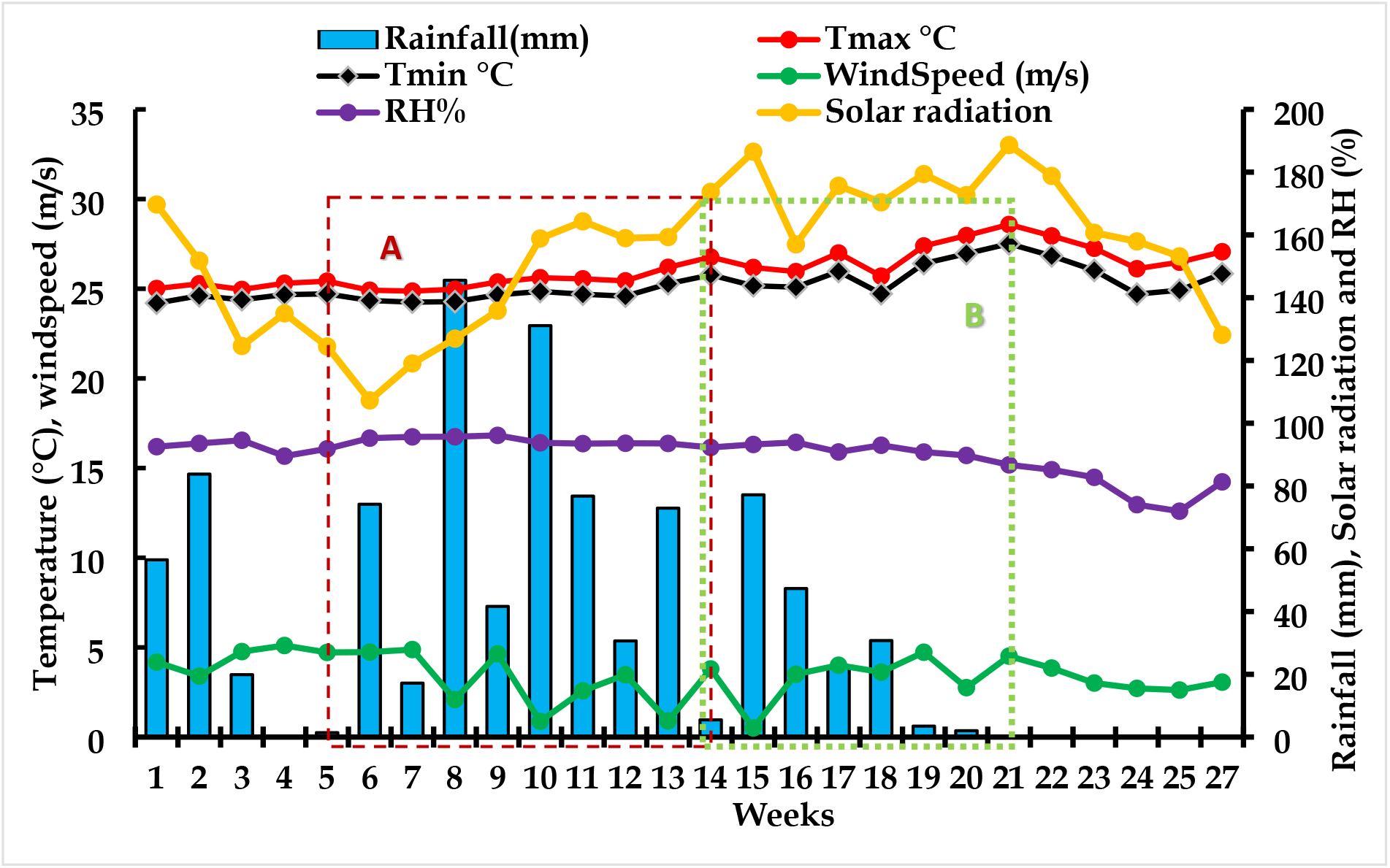
Weekly weather data information of 2020 and 2021 crossing windows, IITA Ibadan station. Week 1 corresponds to 1^st^ week of July while Week 27 corresponds to the last week of December. (**A**) *D. rotundata* crossing window started early August (Week 5) and ended mid-October (Week 14). (**B**) *D. alata* crossing window started in mid-October (Week 14) and ended early December (Week 21).

Among weather conditions, the solar radiation fluctuated much across flowering weeks (107.3 MJ m^−2^ day^−1^ in week 6 to 188.6 MJ m^−2^ day^−1^ in week 21). The wind speed varied with weeks (week 15 had lowest wind speed: 0.5 km ha^−1^ while week 4 had highest speed: 5.1 km ha^−1^). Maximum and minimum temperatures followed similar trends across weeks: week 21 had the highest minimum (27.5°C) and maximum temperatures (28.6°C). The relative humidity ranged from 72.0 to 96.1% (Fig 1).

Weather conditions varied with hours of the day: solar radiation, maximum and minimum temperatures were highest at the midday (11 a.m.–3 p.m.). The relative humidity had an opposite trend than temperatures, midday (1–4 p.m.) having lowest values (63.3–65.5%) than morning and night hours (Fig 2).

**Fig 2.**
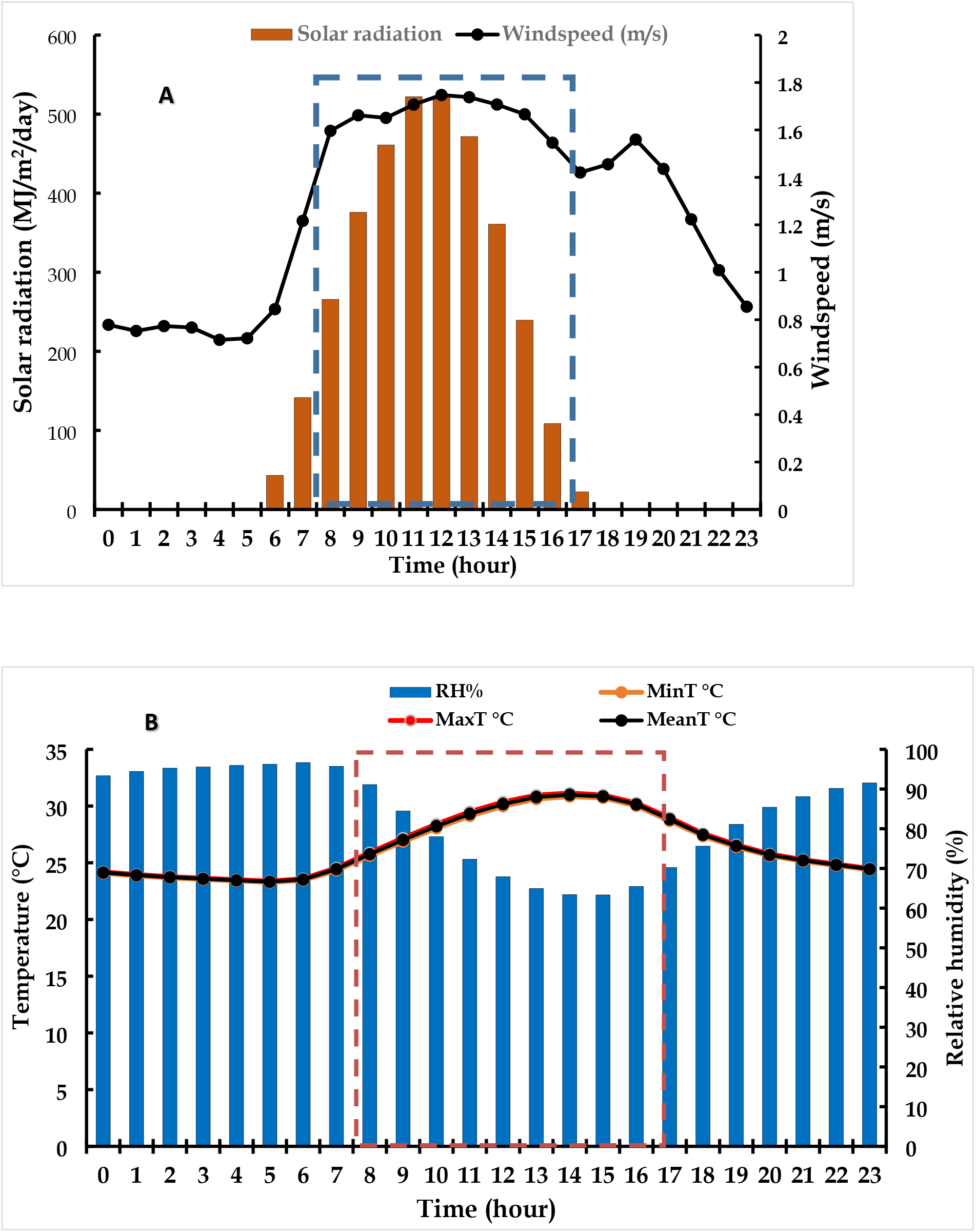
Variations in weather conditions throughout the day during 2020-2021 flowering windows at UTA, Ibadan, Nigeria. On the time axis, 0 refers to midnight while 23 refers to 11 p.m. of the same day. Highlighted hours (8 a.m. to 5 p.m.) correspond to crossing hours. (A) trends of daily variations in solar radiation and windspeed, (B) trends of daily variations in temperatures and relative humidity.

### Pollen viability assessment

*In vitro* pollen germination testing was performed regularly to ensure pollination results were not influenced by the pollen viability status. The previously optimized pollen germination testing protocol by Mondo et al. (2021a) was used. This consisted of culturing anthers with pollen on Petri dishes containing a nutritive medium made of 10% sucrose, 100 ppm H_3_BO_3_, 300 ppm Ca(NO_3_)_2_.4H_2_O, 200 ppm MgSO_4_.7H_2_O, and 100 ppm KNO_3_. This medium was supplemented with 0.5% agar and adjusted at pH 6.5. The culture was incubated at dark for 3 h under 25°C. The pollen germination output was visualized under a fluorescence microscope (Olympus BX51, Tokyo, Japan) at 10× magnification. The stigma receptivity was determined using visual observation of the female flowers prior the crossing.

### Hand pollinations

At flowering, female flowers were bagged with thrip-proof cloth-bags five days before pollination. Hand pollinations between selected male and female plants were carried out from 8:00 a.m. to 5:00 p.m. for the entire flowering window. A cumulative number of 9,775 *D. rotundata* and 6,565 *D. alata* female flowers were hand-pollinated with fresh pollen across crossing hours for the two seasons. At each pollination day, an equal number of female flowers were pollinated hourly. The pollinated flowers were then kept bagged for two weeks to ensure the purity of offspring from crosses.

Following data were collected to assess the optimal time for hand pollination: (1) date of pollination, (2) time of pollination, (3) fruit set (evaluated two weeks after pollination), and (4) the seed set at plant physiological maturity. After fruit processing, the seed viability was also assessed.

Data collected on the fruit and seed sets were further used to calculate the pollination success rate, the seed setting rate (SSR), the seed production efficiency (SPE), and the seed viability (SV) as in Mondo et al. (2021b, 2022). The pollination success rate was calculated as follows:

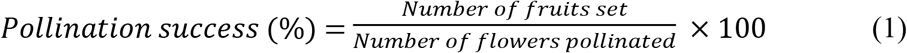

The seed setting rate (SSR) was the ratio between the number of seeds from a cross and the number of fruits multiplied by six (which is the expected number of seeds in a yam fruit):

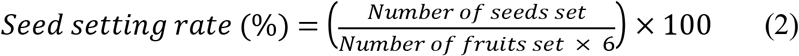

The seed production efficiency (SPE) for a cross was calculated as the number of viable seeds divided by six times the number of pollinated flowers multiplied by 100:

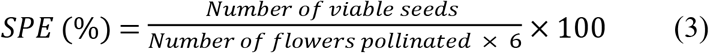

The seed germination rate was estimated by dividing the seedling stand count in nurseries by the number of seeds sown multiplied by 100:

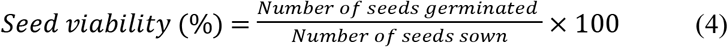

### Statistical data analysis

The analysis of variance (ANOVA) was performed to detect differences among pollination successes, SPE, SSR, and SV using the pollination month, week, and hour of the day as factors. When necessary, means were separated by the least significant difference (LSD) test at 0.05 p-value threshold. Analyses and plotting of graphs were performed using ggplot2 package in R.

## Results

### Pollination success across crossing time

There were significant differences among crossing hours for pollination success in *D. alata* (p = 0.003), morning hours (8–11 a.m.) being better than afternoons (12–4 p.m.) (Fig 3A, S2 Table). No significant difference existed between crossing hours in *D. rotundata* crossing blocks (p = 0.618, Fig 3B) though the mid-day seemed optimal for hand pollination. Based on the crossing block data, 11 a.m. could be considered as optimal for both species (18.6% for *D. alata* and 40.3% for *D. rotundata).* Lowest rates were recorded at 4 p.m. for both species (3.3% for *D. alata* and 30.5% for *D. rotundata).* Though both crossing time and genotype had a significant effect on the pollination outcome (p<0.001), their interaction was not significant (S2 Table). *In vitro* pollen germination data showed, however, that mid-day pollen collection had better results than both extremes though the response was genotype-specific (p = 0.0014; Fig 4, S1 Fig). Pollen geminated most between 12 noon and 2 p.m. (18.7–20% for *D. alata* and 22.9–25.3% for *D. rotundata*).

**Fig 3.**
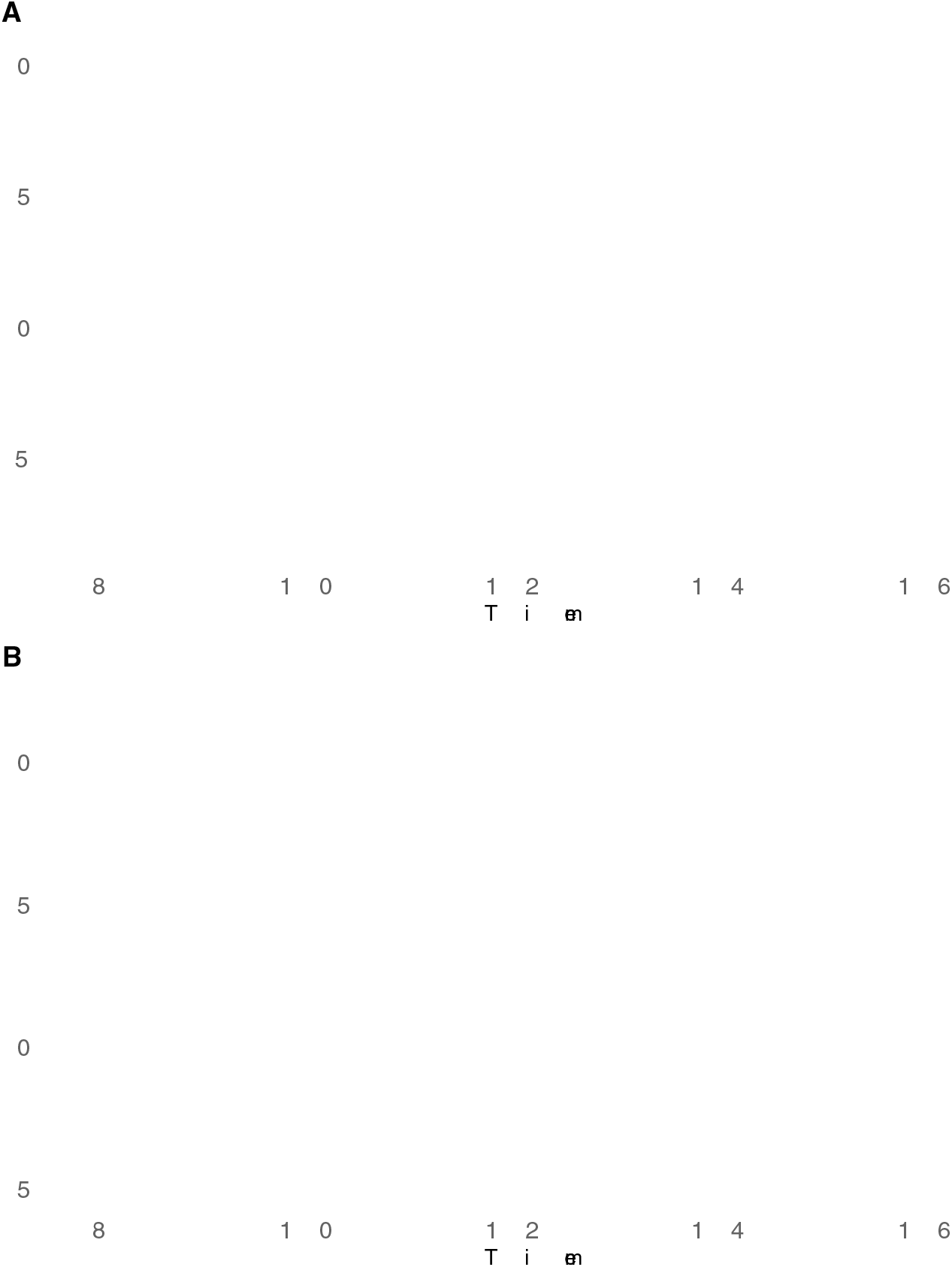
Pollination success across crossing hours: (A) *D. alata,* (B) *D. rotundata*.

**Fig 4.**
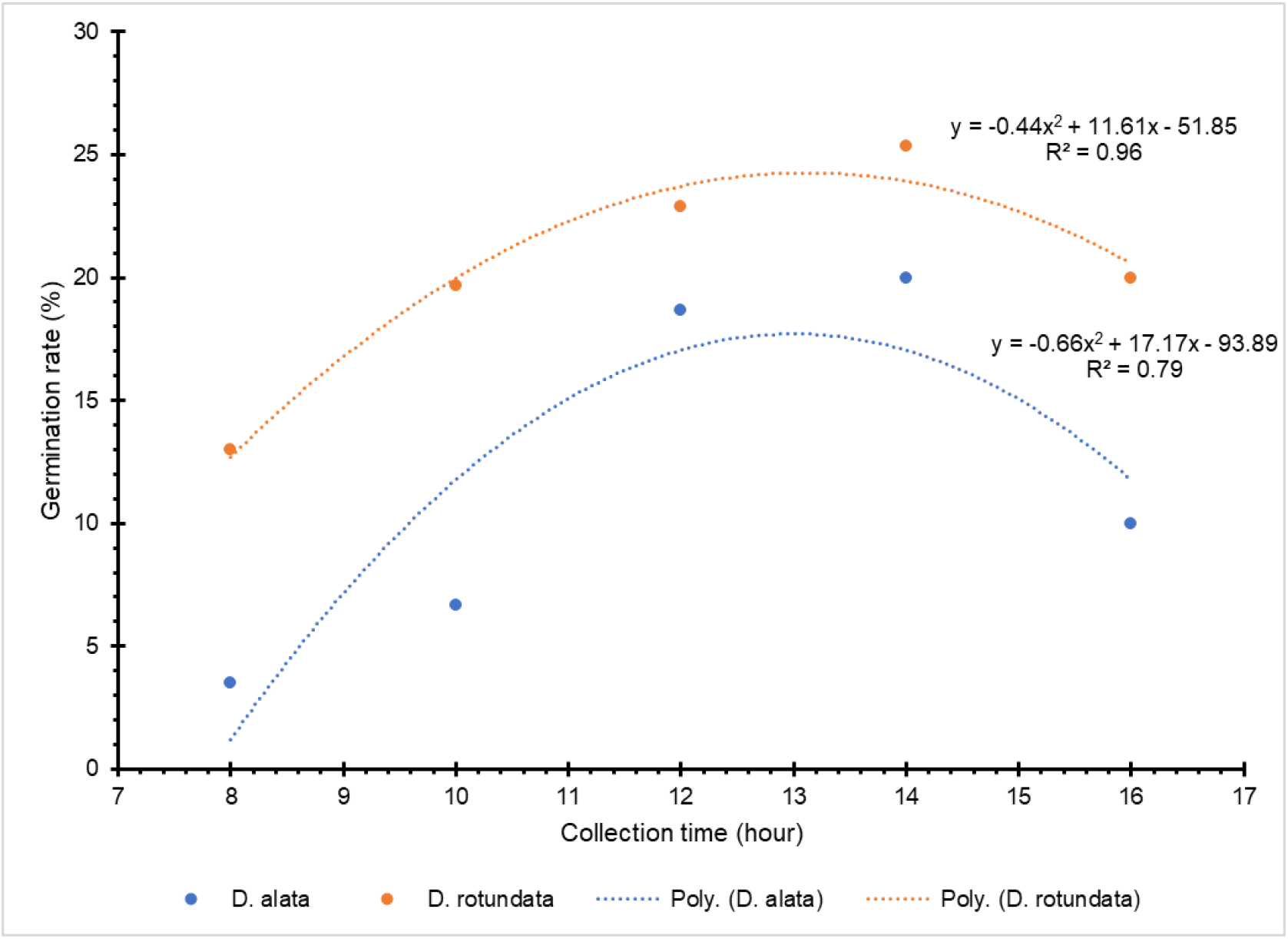
Dynamics of pollen germination rates across day hours.

### Year, month and weekly effects on pollination success and seed set

Pollination success rate was higher in 2021 (29.1%) than 2020 (20.5%) regardless of the species (Fig 5). There were significant differences in pollination success rates among months for *D. alata* (p < 2.2e-16) but not for *D. rotundata* (p = 0.053). During *D. alata* crossing window, October (32.9%) was consistently the optimum month for pollination across both years while November (6.9%) and December (5.6%) had poor pollination success rates. For *D. rotundata* on the other hand, September had relatively best results across years while rates were lowest in October. The pollination success rate varied across weeks within the flowering window (p < 2.2e-16 for *D. alata* and p = 0.004 for *D. rotundata)* (Fig 6). The peak for *D. alata* (35.5–51.0%) was reached during the third week and then started decreasing. *D. rotundata* peak (36.6–68.9%) was observed at week 5 and then started decreasing to reach its minimum pollination rate at week 8 (15.3–19.5%).

**Fig 5.**
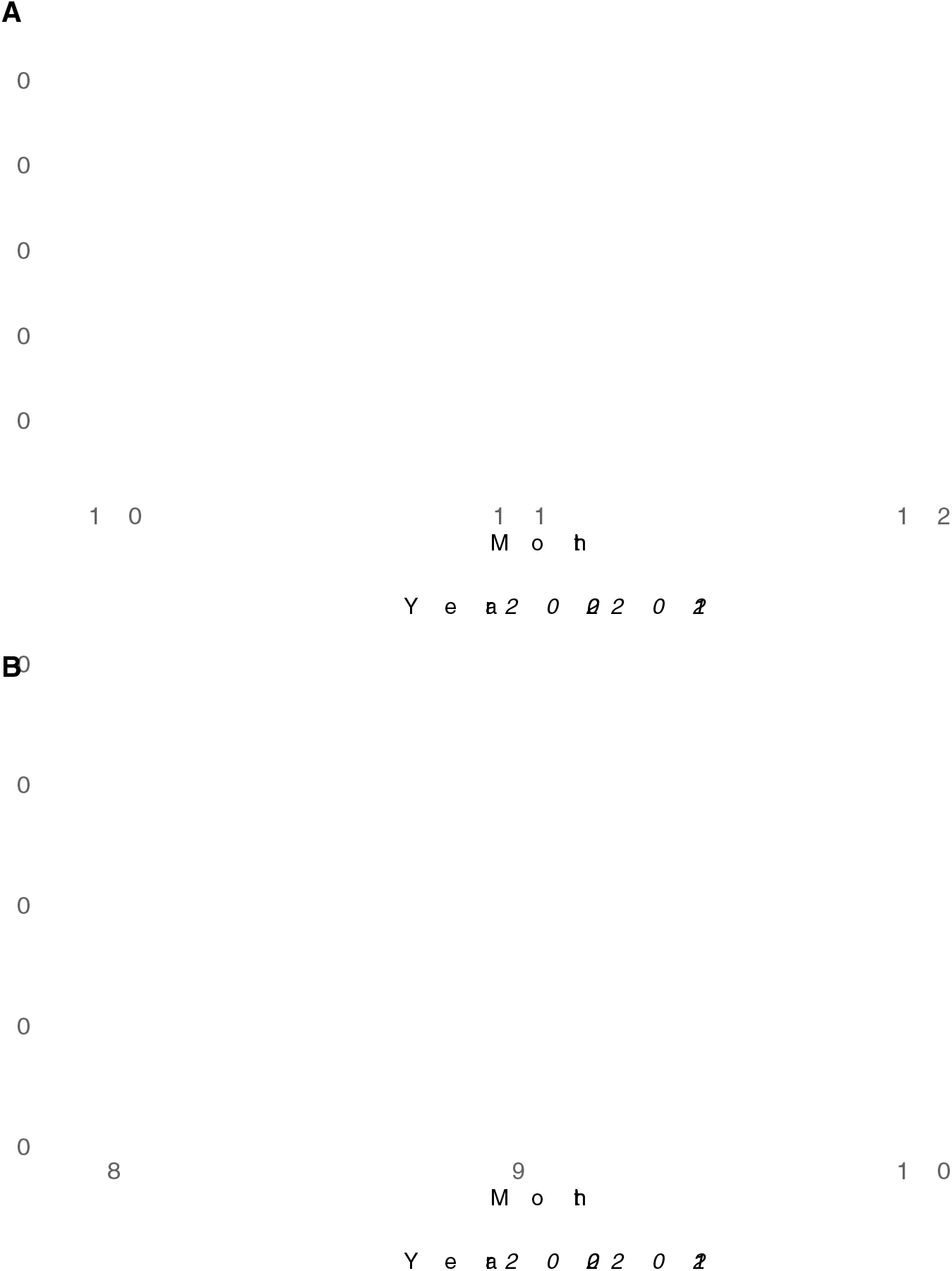
Pollination success rates across crossing months for 2020 and 2021. (A) *D. alata* and (B) *D. rotundata*. The number 8 refers to August and 12 to December.

**Fig 6.**
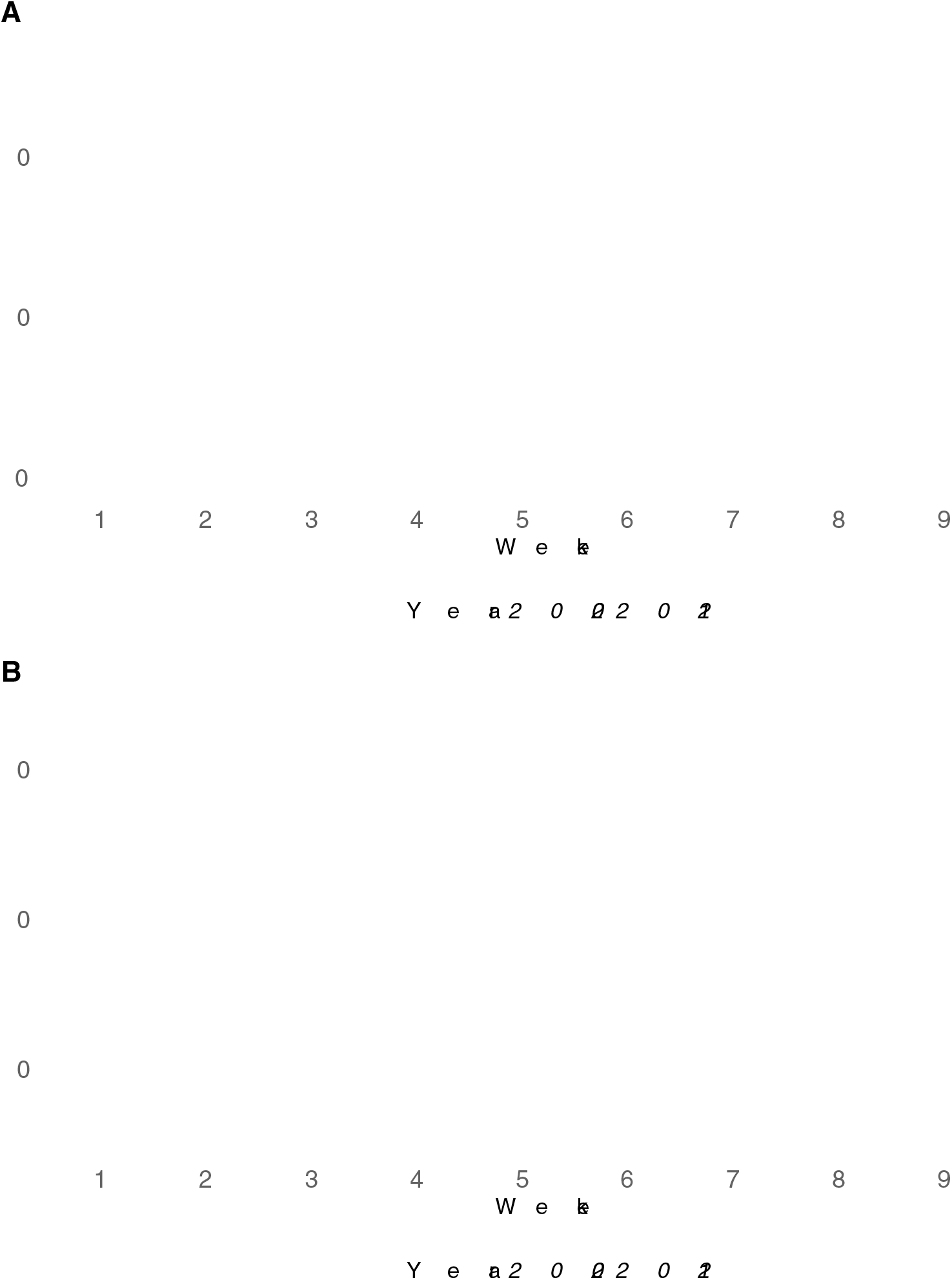
Pollination success rates across crossing weeks for 2020 and 2021. (A) *D. alata* and (B) *D. rotundata.* Week 1 for *D. alata* corresponded to the second week of October while week 9 was the second week of December. Week 1 for *D. rotundata* referred to second week of August while week 9 corresponded to the second week of October.

### Bagging-to-crossing time interval and pollination success

There was an association between the pollination success rate and the bagging-to-crossing time interval (R^2^=0.97, p<0.001). Results suggested that 4–6 days is the optimal interval between flower bagging and crossing time (Fig 7, S2 Fig). The pollination success within that interval ranged from 44.4–50.2% while the lowest success rate was recorded for crosses made within two days after bagging (14.5%).

**Fig 7.**
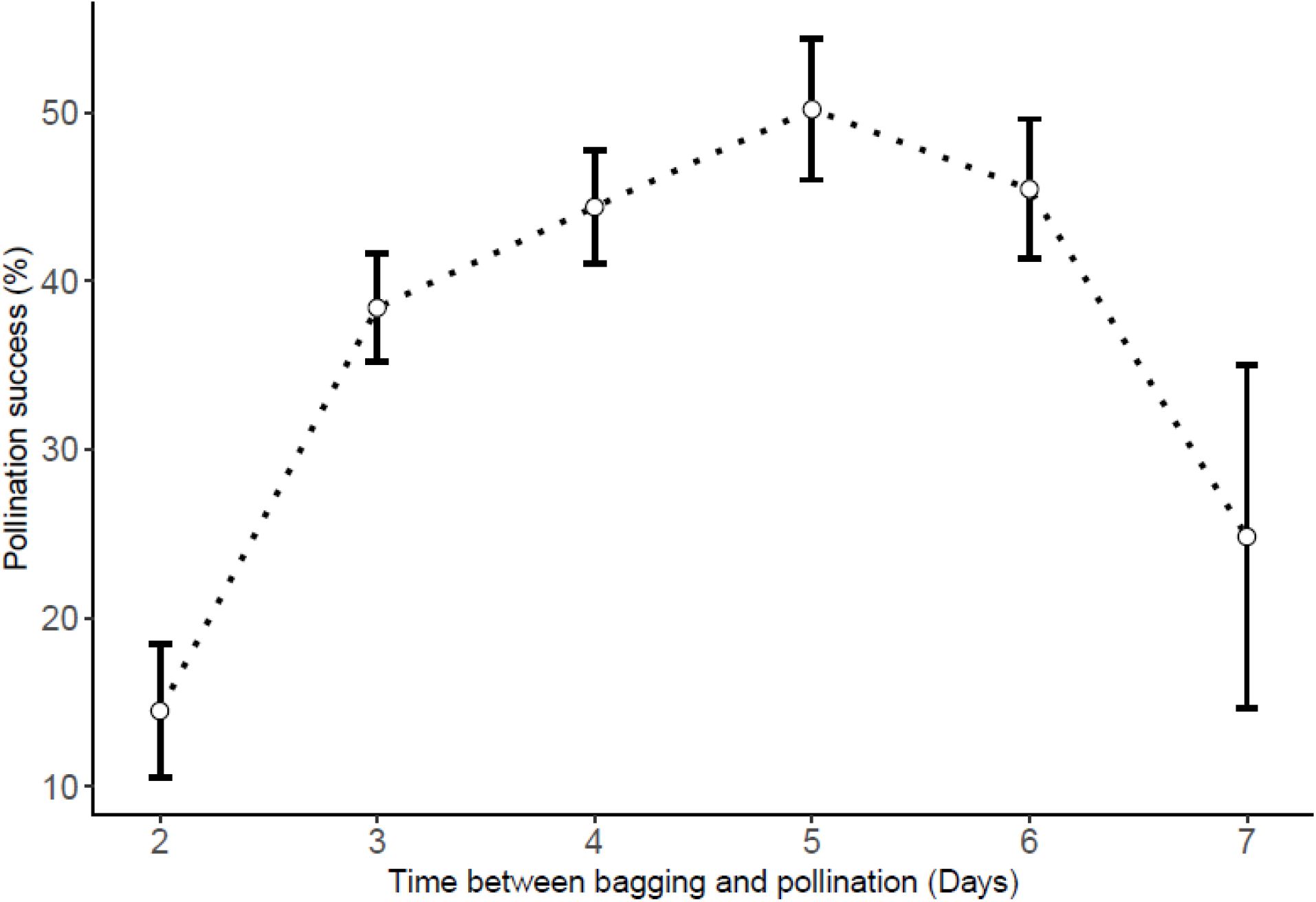
Influence of the time interval between flower bagging and crossing on pollination success rates.

### Dynamics of SPE, SSR, and SV across crossing times

Only the SPE was influenced by the hour of pollination, no particular pattern existed for SSR and SV regardless of the species (Table 1). For SPE, the trend was comparable to the one of the pollination success rate: 11–12 a.m. had highest SPE values (6.01% for *D. alata* and 22.97% for *D. rotundata).* Lowest values were recorded at 4–5 p.m. for both *D. alata* (0.11%) and *D. rotundata* (9.51%). At the monthly basis, the SPE varied with months, October being optimal for *D. alata* (10.3%) and September for *D. rotundata* (18.1%). October (12.2%) and December (0.2%) were worst for *D. rotundata* and *D. alata,* respectively (S3 Fig). For *D. alata,* October had once again the highest SSR (28.1%) and December the lowest (2.3%). August had highest SSR (60.5%) for *D. rotundata* and October the lowest (30.1%) (S4 Fig). Seed viability was indifferent to monthly variabilities and ranged from 70.6–75.6% for *D. alata* and 75.3–81.6% for *D. rotundata* (S5 Fig).

At weekly basis, there were significant differences among weekly SPE (Fig 8), SSR (Fig 9), and SV (Fig 10). Second (10.8) and third (11.8%) weeks had highest SPE for *D. alata* while highest SPE were on fourth (21.9%) and fifth (22.8%) weeks for *D. rotundata.* Lowest SPE were in the 6^th^ to 8^th^ weeks (0%) for *D. alata* and second (7.2%) and 8^th^ (9.9%) weeks for *D. rotundata.* SSR values were not significantly different for weeks 1 to 5 (17.1–29.0%) after which it decreased significantly in *D. alata* crossing blocks (0.0–8.3%). For *D. rotundata,* SSR had a similar trend as for SPE, the second week having lowest SSR (23.3%) and week 1 (65.3), week 4 (60.5%) and week 5 (63.1%) had the highest SSR. *D. alata* seed viability did not vary much across weeks (Fig 10) while for *D. rotundata,* the second week (56.8%) had significantly lower seed viability than all other weekly means (74.8–82.3%).

**Fig 8.**
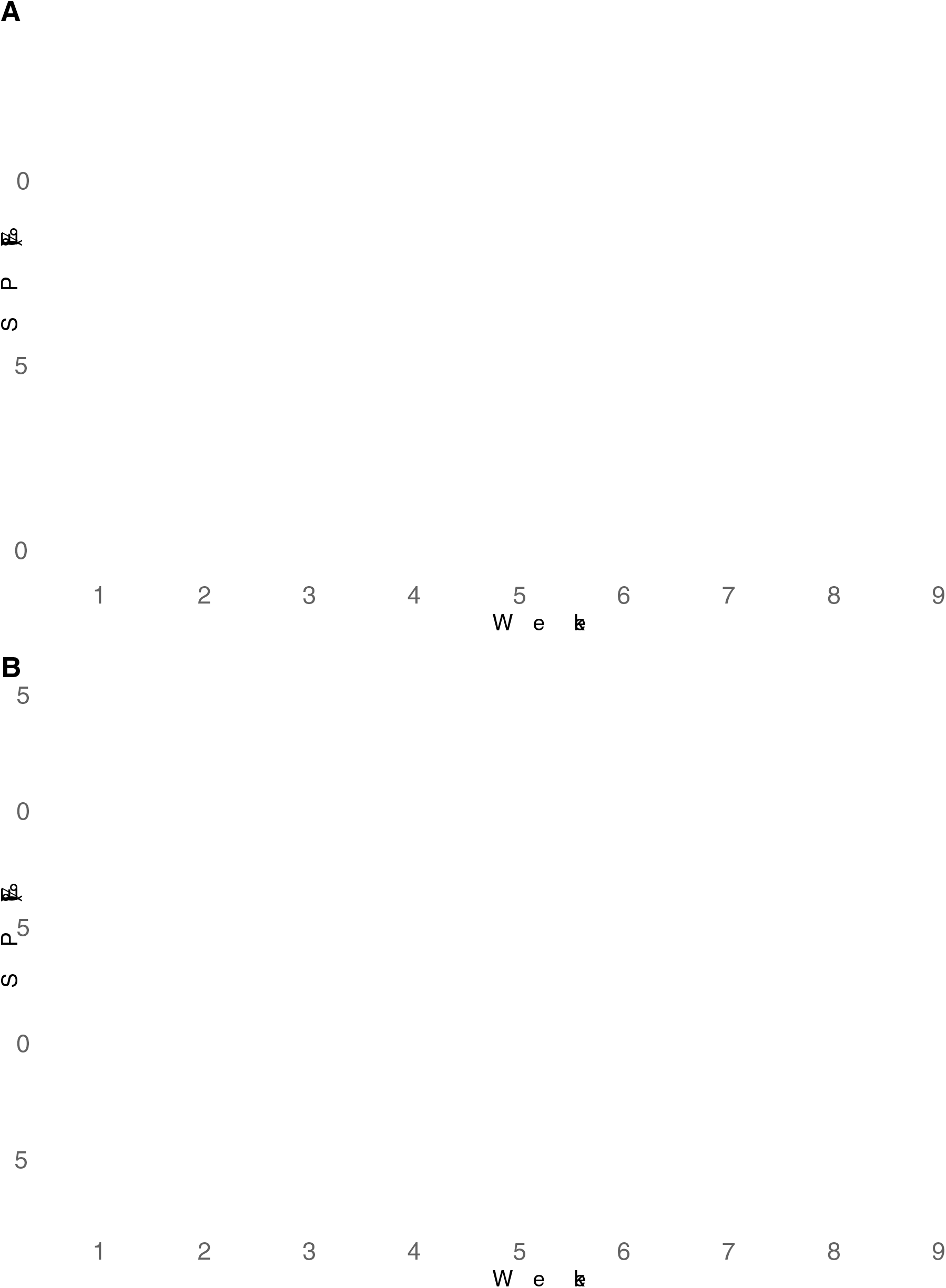
Weekly seed production efficiency across the crossing window: (A) *D. alata* and (B) *D. rotundata.* Week 1 for *D. alata* corresponded to the second week of October while week 9 was the second week of December. Week 1 for *D. rotundata* referred to second week of August while week 9 corresponded to the second week of October.

**Fig 9.**
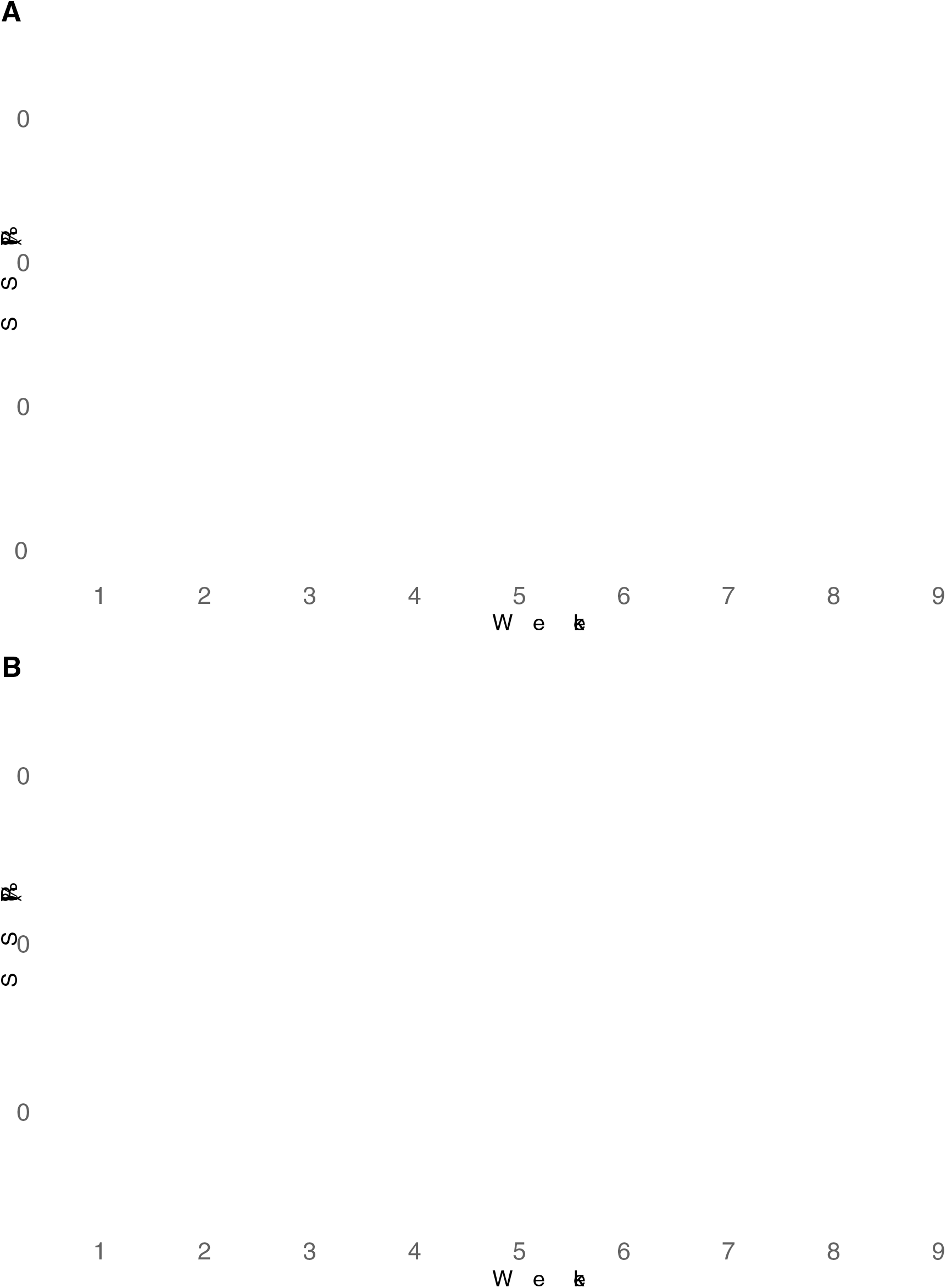
Weekly seed setting rate (SSR) across the crossing windows: (A) *D. alata* and (B) *D. rotundata.* Week 1 for *D. alata* corresponded to the second week of October while week 9 was the second week of December. Week 1 for *D. rotundata* referred to second week of August while week 9 corresponded to the second week of October.

**Fig 10.**
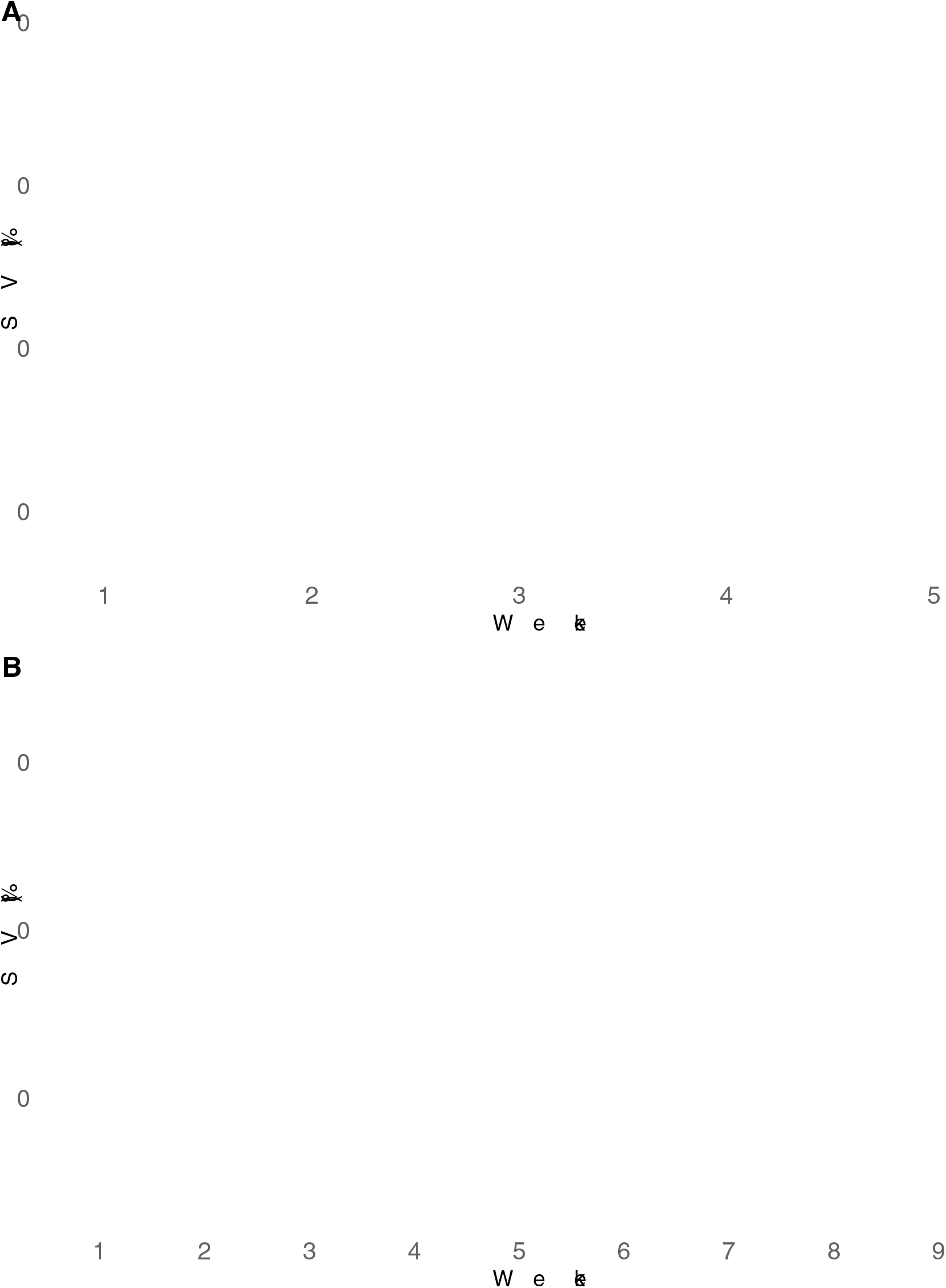
Weekly seed viability rate across the crossing windows: (A) *D. alata* and (B) *D. rotundata.* Week 1 for *D. alata* corresponded to the second week of October. We had no seed for viability test for weeks 6–9. Week 1 for *D. rotundata* referred to second week of August while week 9 corresponded to the second week of October.

**Table 1.**
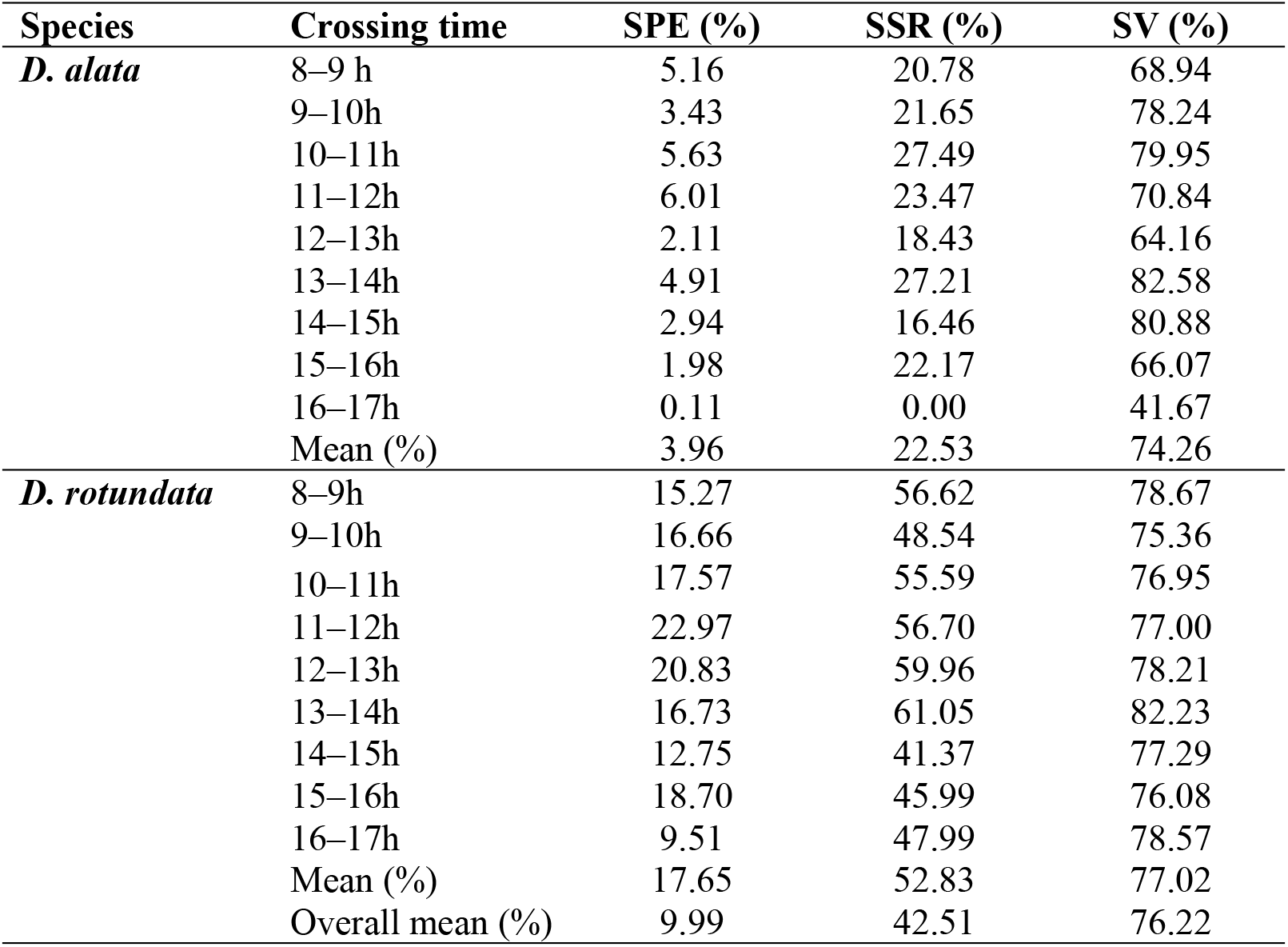
Pollination time and seed production efficiency (SPE), seed setting rate (SSR) and seed viability (SV)

| Species | Crossing time | SPE (%) | SSR (%) | SV (%) |
| --- | --- | --- | --- | --- |
| *D. alata* | 8–9 h | 5.16 | 20.78 | 68.94 |
|  | 9–10h | 3.43 | 21.65 | 78.24 |
|  | 10–11h | 5.63 | 27.49 | 79.95 |
|  | 11–12h | 6.01 | 23.47 | 70.84 |
|  | 12–13h | 2.11 | 18.43 | 64.16 |
|  | 13–14h | 4.91 | 27.21 | 82.58 |
|  | 14–15h | 2.94 | 16.46 | 80.88 |
|  | 15–16h | 1.98 | 22.17 | 66.07 |
|  | 16–17h | 0.11 | 0.00 | 41.67 |
|  | Mean (%) | 3.96 | 22.53 | 74.26 |
| *D. rotundata* | 8–9h | 15.27 | 56.62 | 78.67 |
|  | 9–10h | 16.66 | 48.54 | 75.36 |
|  | 10–11h | 17.57 | 55.59 | 76.95 |
|  | 11–12h | 22.97 | 56.70 | 77.00 |
|  | 12–13h | 20.83 | 59.96 | 78.21 |
|  | 13–14h | 16.73 | 61.05 | 82.23 |
|  | 14–15h | 12.75 | 41.37 | 77.29 |
|  | 15–16h | 18.70 | 45.99 | 76.08 |
|  | 16–17h | 9.51 | 47.99 | 78.57 |
|  | Mean (%) | 17.65 | 52.83 | 77.02 |
|  | Overall mean (%) | 9.99 | 42.51 | 76.22 |

## Discussion

### Pollination success depended on crossing time

Yam breeding is challenged by sexual reproduction abnormalities (sparse, irregular, and asynchronous flowering, cross compatibility barriers and low fertility) because of the domestication process that favored vegetative propagation at the expense of botanical seeds (Mondo et al., 2020). The International Institute of Tropical Agriculture (IITA) has devised a series of studies to control those reproduction abnormalities. This study is a continuation of such efforts and aimed to update recommendations on optimum time for hand pollination in yam breeding. We realized that the pollination success varied with the year, month, week, and hour of the day when the hand pollination is performed. The year, month, week, and hour of the day with well-distributed rainfall, high relative humidity and moderate temperatures were conducive to hand pollination for both *D. alata* and *D. rotundata* yam species. The year 2021 was more wet than 2020 and thus recorded higher pollination success rate than 2020. That finding agreed with Mondo et al. (2022) who showed variability in pollination success across years while using 2010–2020 crossing block information. *D. alata* was more sensitive to the time of pollination than *D. rotundata.* For instance, Mondo et al. (2021a) showed that *D. rotundata* pollen had a wide range of germination temperatures (15–35°C) compared to *D. alata* that gave better results at 25°C. The weather parameters’ fluctuation with time could have explained the difference in pollination outcomes. A previous study on yam showed that weekly variability in rainfall, temperature, relative humidity, sunshine, and the number of rainy days within the yam flowering window (July to November) significantly influenced the pollination outcomes in either the *D. rotundata* or *D. alata* crossing blocks (Mondo et al., 2022). Abraham and Nair (1990) also reported that successful pollination in yam is significantly associated with high relative humidity and moderate atmospheric temperatures. As supported by this study, making crosses in some weeks of August or in the second half of October for *D. rotundata* and after 15^th^ November for *D. alata* could result in low pollination success due to the suboptimal weather conditions. As recommended by Mondo et al. (2022) there is the need for supplemental irrigation in yam crossing blocks to reduce the water deficit’s adverse effects on yam reproductive phases during these months. Since *D. alata* presents no dormancy at harvest, options of establishing the crossing block as early as March (with supplemental irrigation) could help avoiding the coincidence of its flowering window with harsh environmental conditions, as it was the case the last two years.

In contrast to most reports on yam pollination, *D. rotundata* had higher crossability rate and higher seed production efficiency than *D. alata* for both 2020 and 2021. However, *D. alata* had consistently higher values (31%) than *D. rotundata* (23%) when bulking 2010–2020 crossing block information at IITA (Mondo et al., 2022). For our study period (2020–2021), months corresponding with the *D. alata* flowering window (October–December) were drier compared to those of *D. rotundata* (August–October) which benefited from relatively high and well-distributed rains and moderate temperatures. It is noteworthy that *D. alata* is sensitive to rainfall distribution, sunshine, relative humidity, and temperatures (Abraham and Nair, 1990; Mondo et al., 2022) which were suboptimal during *D. alata* flowering window.

For both seasons, crosses made in morning hours (8–12 a.m.) had better results than those from afternoons for *D. alata*. Though not significant, mid-day seemed optimal for *D. rotundata*. Based on the crossing block data, 11 p.m. could be considered as optimal for both species while lowest rates were recorded at 4 p.m. for both species. However, *in vitro* pollen germination tests supported the mid-day (12 noon–2 p.m.) as the optimal time for pollen collection for both species while both morning and evening extremes should be avoided. This result aligned with findings by Akoroda (1983, 1985) that showed that *D. rotundata*’s better pollination success was achieved when crosses are made between 12 noon and 2 p.m. at IITA Ibadan, Nigeria. These findings partly dismissed our hypothesis that the optimal time for hand pollination, recommended four decades ago, might have been affected by climate changes. Since weather conditions are conducive for human labor in morning hours than the mid-day, we could recommend concentrating crossing activities at 11–12 a.m. interval for both species since morning was better than afternoons for *D. alata* and there were no significant differences between morning and mid-day hours for *D. rotundata*. Results showed an influence of the flower bagging on pollination outcomes, 4–6 days being the optimal interval between flower bagging and crossing time. Further investigations are necessary to elucidate reasons behind the influence of bagging-to-crossing time interval in yam crossing blocks.

#### The seed setting rate and the seed viability were less affected by the hour of crossing but varied with the month and week of crossing

Only the pollination success rate and the SPE varied with the hour of pollination, no particular pattern existed for the SSR and SV regardless of the species. For the SPE, the trend was comparable to the one of the pollination success rate: 11–12 a.m. had highest SPE values for both species while lowest values were recorded at 4–5 p.m. At the monthly basis, the SPE varied with months, October being optimal for *D. alata* and September for *D. rotundata.* October and December were worst for *D. rotundata* and *D. alata*, respectively. For *D. alata*, October had once again the highest SSR and December the lowest. August had highest SSR for *D. rotundata* and October the lowest. Seed viability was indifferent to monthly variabilities for both species.

There were significant differences among weekly SPE, SSR, and SV, weeks with conducive climatic conditions provided the best outcomes. Bandeira e Sousa et al. (2021) showed that environmental factors (temperature, rainfall, and photoperiod) contribute to a post-zygotic barrier in crops like cassava. They showed that high temperatures induced flower abortion and reduced the number of female flowers per inflorescence and seed setting rate. There was also a decreased pollen tube growth rate at higher average temperatures than lower temperatures, supporting the hypothesis that environmental conditions affect the efficiency of sexual reproduction, and that appropriate planning of planting dates and locations can maximize seed production (Ramos Abril et al., 2019; Bandeira e Sousa et al., 2021). Environmental factors such as rainfall and temperature had also affected flowering, pollen production, and fruit development in cocoa (Omolaja et al., 2009).

### Conclusion

This study, based on two-year crossing data, showed that the time of pollination had an influence on the pollination success rates. The year, month, week, and hour of the day with well-distributed rainfall, high relative humidity and moderate temperatures were conducive to hand pollination for both yam species. Crossing block data, weather information, and *in vitro* pollen germination seemed to encourage morning to mid-day hybridization for better pollination results. Special measures should be devised for *D. alata* as it was the most sensitive to weather conditions, and the months corresponding to its flowering window had globally suboptimal climatic conditions for both years.

## Supporting information

**S1 Fig. Genotypic variability in pollen germination in *D. alata* and *D. rotundata***

**S2 Fig. Relationship between the bagging–crossing time interval and pollination success**

**S3 Fig. Monthly variability in SPE for *D. alata* and *D. rotundata***

**S4 Fig. Monthly variability in SSR for *D. alata* and *D. rotundata***

**S5 Fig. Monthly variability in seed viability for *D. alata* and *D. rotundata***

**S1 Table. Description of yam genotypes used in hand pollination experiment**

**S2 Table. ANOVA table for pollination success across crossing hours for *D. alata***

## Acknowledgments

The funding support from the Bill & Melinda Gates Foundation (BMGF) is acknowledged. We are also grateful to the yam breeding team at IITA Ibadan for active participation in data collection. The first author is grateful for the African Union Commission’s scholarship for his Ph.D. studies at the Pan African University-Institute of Life and Earth Sciences (PAULESI). Yannick Mugumaarhahama and Géant B. Chuma are acknowledged for statistical advice.

## Data Availability Statement

All relevant data are within the manuscript and its Supporting Information files.

## Author Contributions

**Conceptualization:** Asrat Asfaw, Jean M. Mondo, Paterne A. Agre

**Data curation:** Jean M. Mondo

**Formal analysis:** Jean M. Mondo, Paterne A. Agre

**Funding acquisition:** Asrat Asfaw

**Investigation:** Jean M. Mondo

**Methodology:** Asrat Asfaw, Jean M. Mondo, Paterne A. Agre

**Project administration:** Asrat Asfaw, Paterne A. Agre

**Resources:** Asrat Asfaw

**Supervision:** Asrat Asfaw, Paterne A. Agre, Malachy O. Akoroda

**Writing – original draft:** Jean M. Mondo, Paterne A. Agre, Asrat Asfaw

**Writing – review & editing:** All authors

